# D2/D3 dopamine supports the precision of mental state inferences and self-relevance of joint social outcomes

**DOI:** 10.1101/2023.05.02.539031

**Authors:** J.M. Barnby, V. Bell, Q Deeley, M. Mehta, M. Moutoussis

## Abstract

Striatal dopamine is important in paranoid attributions, although its computational role in social inference remains elusive. We employed a simple game theoretic paradigm and computational model of intentional attributions to investigate the effects of dopamine D2/D3 antagonism on ongoing mental state inference following social outcomes. Haloperidol, compared to placebo, enhanced the impact of partner behaviour on beliefs about the harmful intent of partners, and increased learning from recent encounters. These alterations caused significant changes to model covariation and negative correlations between self-interest and harmful intent attributions. Our findings suggest haloperidol improves belief flexibility about others and simultaneously reduces the self-relevance of social observations. Our results may reflect the role of D2/D3 dopamine in supporting self-relevant mentalisation. Our data and model bridge theory between general and social accounts of value representation. We demonstrate initial evidence for the sensitivity of our model and short social paradigm to drug intervention and clinical dimensions, allowing distinctions between mechanisms that operate across traits and states.

**Data Availability:** All data and code are available online:

https://github.com/josephmbarnby/Barnby_etal_2023_D2D3Modelling

**Graphical Abstract:** 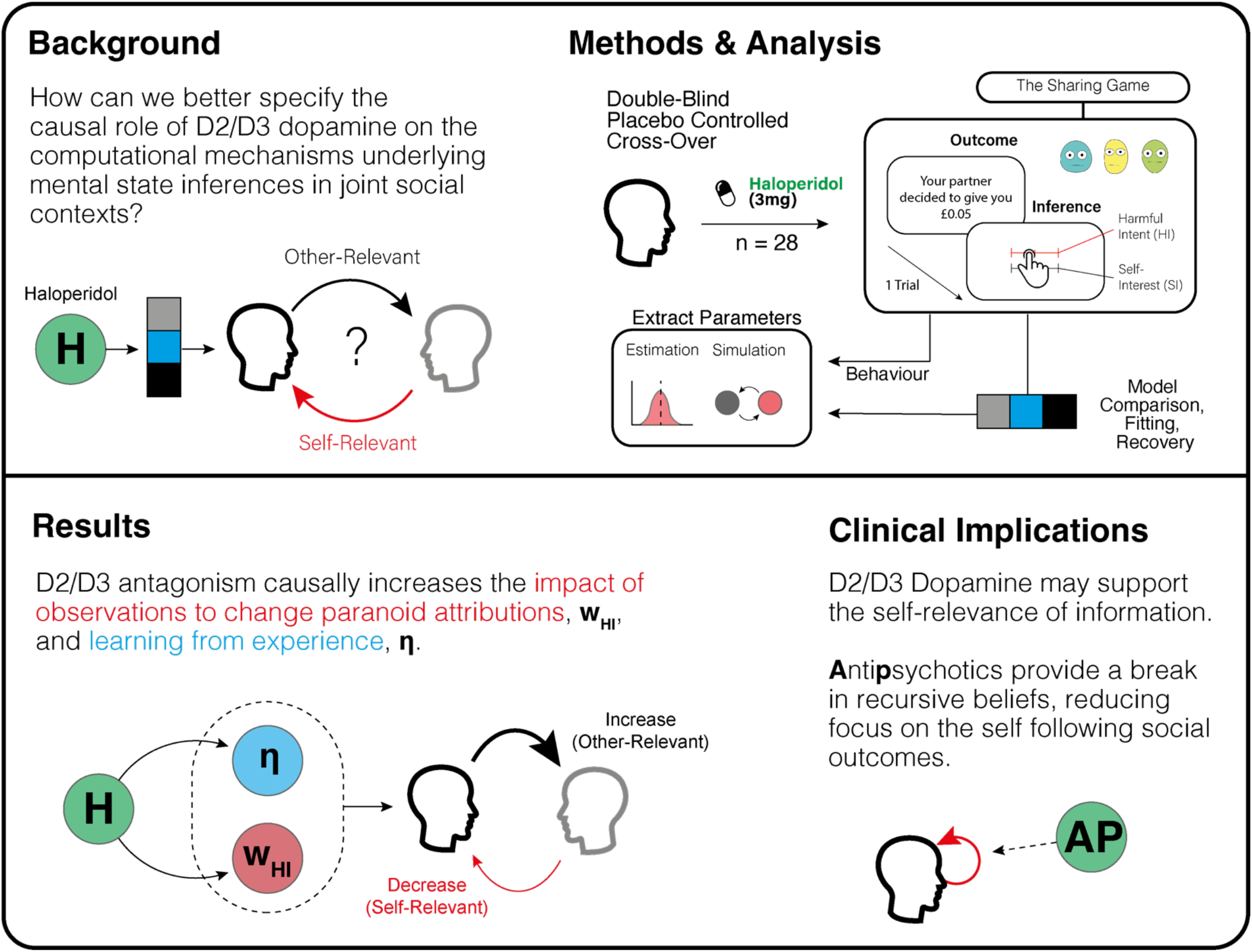

## Introduction

Dysregulated striatal dopamine has been identified as a key causal component in psychosis. Influential work proposed that striatal dopamine mediates aberrant salience leading to atypical perceptual experiences [1-3] more recent social-developmental models have highlighted the role of dopamine as a key point of convergence for a number of causal social and developmental factors, such as trauma, genetic vulnerability, and cannabis use [4]. This has been supported by molecular and neuroimaging studies suggesting that developmental adversities (e.g., [5,6]) increases pre-synaptic turnover of dopamine in striatal regions that may fuel the onset [7-9] and exacerbation [10,11] of psychosis symptoms.

Antipsychotics are the first-line treatment for psychosis and have good evidence for their efficacy [12]. While they are thought to enact their therapeutic efficacy via D2/D3 dopamine antagonism, the exact mechanism by which their pharmacological effect reduces symptoms through the modulation of neurocognitive processes is still poorly understood. Although recent work on the links between striatal hyperdopaminergia and psychosis has been important in identifying important risk factors and has offered important hypotheses for the causes of psychosis and psychotic symptoms at the neurobiological level, it has not been able to explain how they alter cognition beyond citing salience as a key mechanism. The end point of such causal pathways in psychiatry are likely to be dynamic, multi-dimensional, context-sensitive cognitive processes [13]. Computational modelling is an approach that allows these dynamic cognitive processes to be mathematically implemented and has the potential to connect mechanism more effectively to psychiatric phenomenology [14,15], offering precise accounts of complex behaviour that are more amenable to formal testing, refutation and refinement. Within this framework, dopaminergic alterations have been linked to computational processes such as belief updating [16,17], expectations of belief volatility [18-20], and model-based control [21].

One particularly disabling core symptom of psychosis is paranoia, the unfounded belief that others are trying to cause you harm [22,23]. Psychologically, paranoia is characterised by heightened sensitivities to interpersonal threat [24], attributing negative outcomes to external, personal causes [25], and overly complex mentalisation [26-27]. Developing computational theories to bridge the gap between the phenomenology and the neurocognitive mechanisms of paranoia requires particular considerations. Computational approaches in the social domain must sufficiently account for large, and often recursive, action spaces [28]. These structural principles are appropriate for psychiatric symptoms which inherently involve alterations to interpersonal beliefs concerning the self and others [29].

Models of intentional attributions – explicit inferences about the mental state of others - allow for analyses that are theoretically related to ongoing paranoia. Current models include mechanistic explanations for perceived changes in the harmful intent and self-interest that might motivate the actions of another. Prior work suggests high trait paranoia is associated with rigid priors about the harmful intent of partners, and a belief that a partner’s actions are not consistent with their true intentions [30,31]. Several predictions can be made concerning the influence of dopamine D2/D3 antagonism on paranoia. Synthetic, *in silico* models [32], neuroimaging evidence [33], prior predictions [31], and parallel psychopharmacological work [21,34] predict that D2/D3 antagonism will increase belief flexibility and improve consistency of the self’s model of others, which in turn should reduce self-relevant attributions of harmful intent following social outcomes. However, this has yet to be tested.

While key binding sites of most antipsychotics are thought to work through their action at D2/D3 dopamine receptors, how they influence the cognitive processes of paranoia is unknown. Given the experimental evidence and synthetic predictions on the role of D2/D3 dopamine antagonism on improvements in belief updating, reductions in harmful intent, increases in prosocial behaviours, and the impact of high trait paranoia on the consistency of a self’s model of others, it follows that the mechanism of action of D2/D3 antagonism on harmful intent attributions may occur through an increase in belief flexibility and the consistency of a self’s model of others. Following from our preregistered behavioural experiment [35], we further examine the causal influence of D2/D3 dopamine receptor antagonism on computational mechanisms governing intentional attributions within a simple game theoretic context. Using a formal model of intentional attributions and an iterative Dictator game [30,31], we test the impact of haloperidol, a D2/D3 antagonist, and L-DOPA, a presynaptic dopamine potentiator, on paranoid beliefs using past data [35].

Primarily we assessed whether haloperidol alters key computational processes involved in mental state inferences, allowing distinctions between trait representational changes (priors) and state learning processes (policy flexibility, uncertainty) along each attributional dimension (harmful intent and self-interest). Given the absence of any consistent descriptive effects of L-DOPA in this experiment we modelled the data under an assumption that there would be no opposing effects on model parameters under LDOPA vs. haloperidol.

## Methods

### Participants

This study was approved by KCL ethics board (HR-16/17-0603). All data were collected between August 2018 and August 2019. Participants were recruited through adverts in the local area, adverts on social media, in addition to adverts circulated via internal emails.

Eighty-six participants were preliminarily phone screened. 35 participants were given a full medical screen. Thirty healthy males were recruited to take part in the full procedure. Two failed to complete all experimental days, leaving 28 participants for analysis. Inclusion criteria were that participants were healthy males, between the ages of 18 and 55. Participants were excluded if they had any evidence or history of clinically significant medical or psychiatric illness; if their use of prescription or non-prescription drugs was deemed unsuitable by the medical team; if they had any condition that may have inhibited drug absorption (e.g. gastrectomy), a history of harmful alcohol or drug use determined by clinical interview, use of tobacco or nicotine containing products in excess of the equivalent of five cigarettes per day, a positive urine drug screen, or were unwilling or unable to comply with the lifestyle guidelines. Participants were excluded who, in the opinion of the medical team and investigator, had any medical or psychological condition, or social circumstance, which would impair their ability to participate reliably in the study, or who may increase the risk to themselves or others by participating. Some of these criteria were determined through telephone check for non-sensitive information (age, gender, general understanding of the study, and overall health) before their full screening visit.

### Procedure

This study was part of a larger study that assessed the role of dopaminergic modulation on personality, beliefs, and social interaction. Here, we focus on the role of dopamine antagonism and pre-synaptic increases in the attribution of mental state inferences during a Dictator game (described below; see Figure 1a).

**Figure 1.**
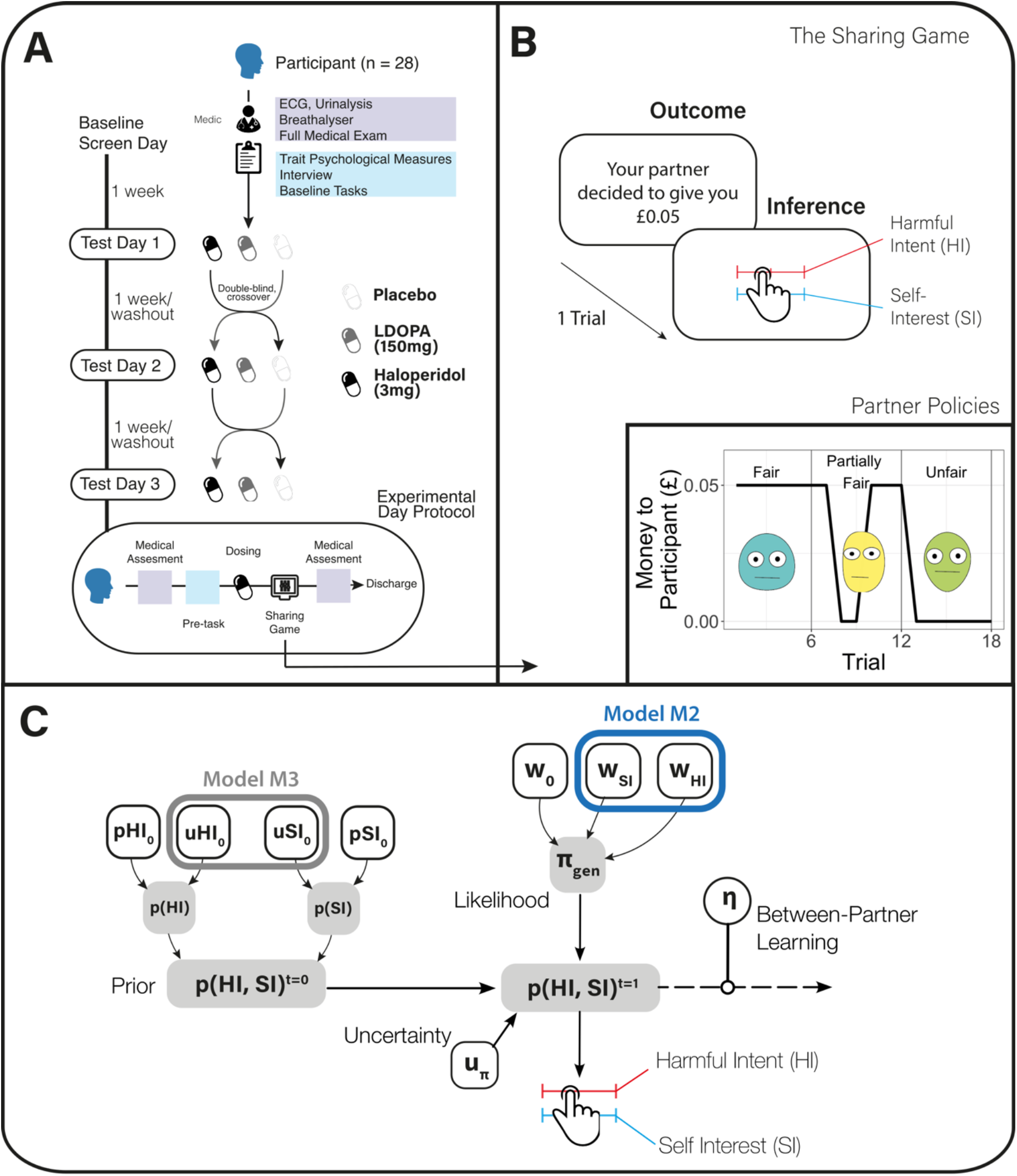
Experimental design and model space. (A) Participants were entered into a double-blind, placebo-controlled, within-subject experimental design. (B) Participants engaged in a three-partner version of the sharing game. Here, partners were assigned the role of Dictator and on each trial could either take £0.10 for themselves (unfair outcome) or take £0.05 and give the participant £0.05 (fair outcome). Participant reported two types of attributional intent concerning the motivations of the partner after each outcome. These included harmful intent attributions and self-interest attributions. Partner order was randomised, and partner change was signalled. (C) Model space used to test whether dopamine manipulations were best explained by the full model (M1), a model that constrained policy updating to a single sensitivity parameter for each attribution (M2), or a model that constrained prior uncertainty to a single parameter (M3; Table 1). White filled objects are free parameters. Grey shaded objects are probability distributions.

The full procedure for participant screening is documented in a prior publication [35]. Briefly, participants who passed the brief phone screening were invited to attend an on-site screening day (see above). Participants were tested for drugs of abuse (SureScreen Diagnostics Ltd) and alcohol (breath test) prior to each experimental day and were excluded if any test was positive. Participants were given at least 7 days, but no more than two months, in between experimental days to allow for drug washout.

On experimental days, participants were randomised to be initially administered either a placebo or 3mg haloperidol in two capsules, and 10mg of domperidone (to reduce known side effects of vomiting and nausea that can appear in some recipients) in one capsule (3 caps total). After half an hour, participants were dosed a second time with either 150mg of co-beneldopa (herein referred to as L-DOPA) or placebo in two capsules. Participants would never receive haloperidol and L-DOPA in the same day.

### The Sharing Game

Participants were asked to play a within-subjects, multi-trial modification on the Dictator game design used in previous studies to assess paranoia [35,36], hereafter called ‘The Sharing Game’ (Figure 1b). In the game, participants played six trials against three different types of partner who are assigned the role of Dictator. In each trial, participants were told that they have a total of £0.10 and their partner (the Dictator) had the choice to take half (£0.05) or all (£0.10) the money from the participant. Partner policies were one of three types: always take half of the money, have a 50:50 chance to take half or all of the money, or always take all of the money. These policies were labelled as fair, partially fair, and unfair, respectively. The order that participants were matched with partners was randomised. Each partner had a corresponding cartoon avatar with a neutral expression to support the notion that each of the six trials was with the same partner.

After each trial, participants were asked to rate on a scale of 1–100 (initialised at 50) to what degree they believed that their partner was motivated (a) by a desire to earn more (self-interest), and (b) by a desire to reduce their bonus in the trial (harmful intent). From the participants perspective, the actions of the partner can be framed as either arising from motivations that concern the gain of value for the partner irrespective of the participant (other-relevant) or arising from motivations that concern the loss of value for the participant (self-relevant).

After making all 36 attributions (two trial attributions for each of the six trials over three partners), participants were put in the role of the Dictator for six trials—whether to make a fair or unfair split of £0.10. Participants were first asked to choose an avatar from nine different cartoon faces before deciding on their six different splits. These Dictator decisions were not used for analysis but were collected to match subsequent participants with decisions from real partners. Participants were paid a baseline payment for their completion, plus any bonus they won from the game.

### Analysis

Behavioural data has been previously published [35]. Here, we apply three computational hypotheses which could explain the data, centred around a Bayesian model [31] developed to explain mental state inference dynamics during social observation, where recursive, strategic social action is not a process of interest [29]. We note that previous work showed a Bayesian instantiation of this attributional model outperformed associative model variants [31]. Model 1 allowed separate uncertainties and likelihood weights for each attribution, identical to our prior work [31]; this model demonstrated that trait paranoia increased belief rigidity and self-other inconsistency, and by extension, may serve as a useful assay to test the mechanisms of haloperidol which is theorised to reduced paranoia. In line with general theories of belief updating [37], Model 2 hypothesised that beliefs would be updating with the same likelihood weight. Model 3 hypothesised that prior beliefs share a single uncertainty free parameter over each distribution, allowing for a simpler hypothesis that prior uncertainties may be represented by a single dimension, giving a more parsimonious account of the data. Descriptions of the parameters within the winning model are in Table 1.

**Table 1.**
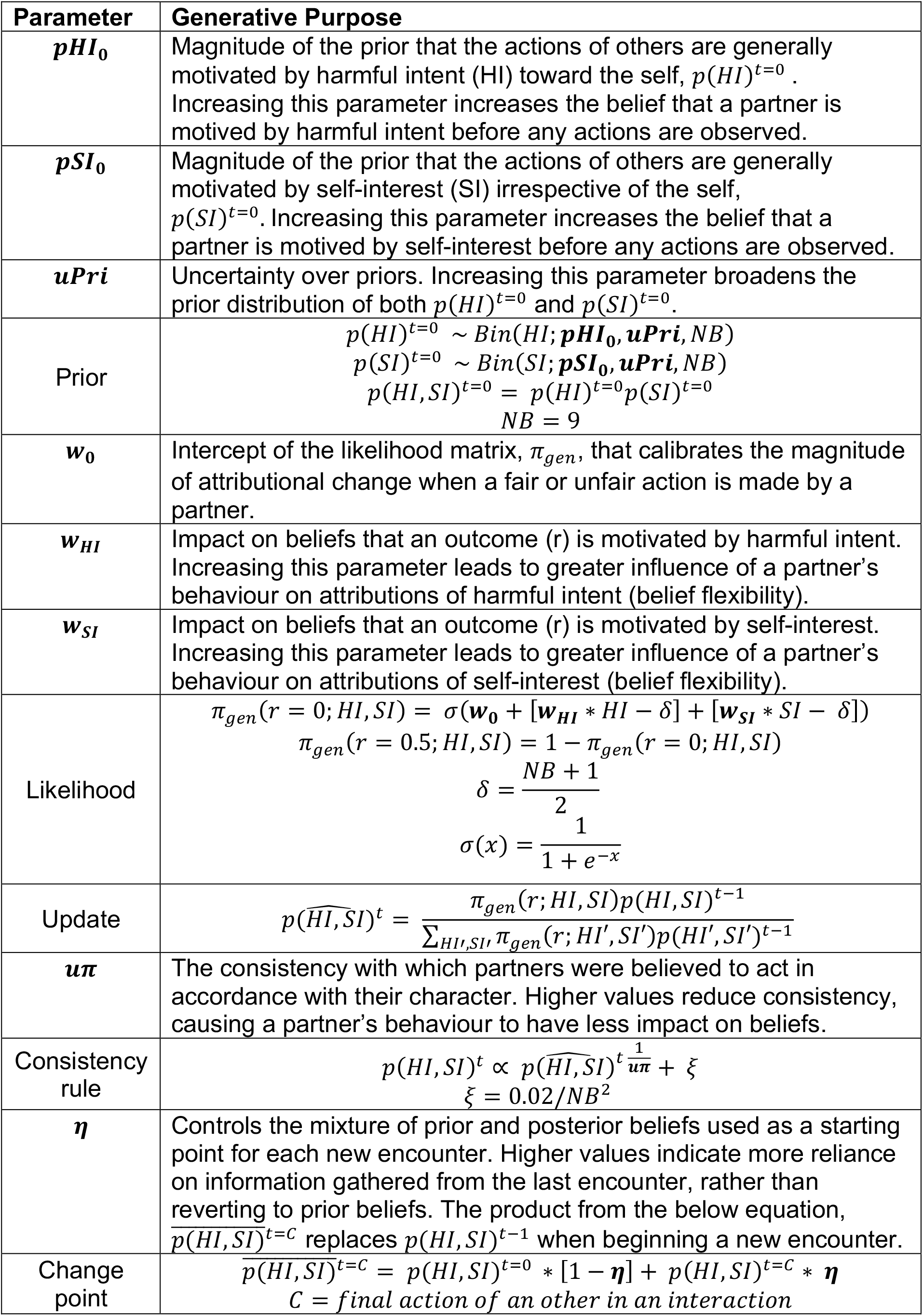
Winning model parameters and their role in the model. By using model fitting procedures modellers can invert the model to approximate the parameter values that may give rise to the observed data. This includes the hidden, prior beliefs of each participant given the variance and magnitude of observed attributions. Using fitted parameter values to simulate each participant allows for generation of pseudo-experimental data - in this case, an agent’s reported intentional attributions, which we can directly compare with the real data. This also approximates the prior beliefs of each participant given the variance and magnitude of observed attributions. *NB* = number of bins discretising the variable represents each attribution; in this case each distribution is comprised of 9 bins. *Bin* = binomial distribution with an added precision parameter, i.e. in the case of HI: *p*(*HI*)^)*t*= 0^ ∼ *Bin*(*HI*; ***pHI***_**0**_, ***uPri***, *NB*)=*p*(*HI*)^)*t*=0^ ∼ *B*(*HI*; ***pHI***_**0**_, *NB*)^1/***uPri***^.

| Parameter | Generative Purpose |
| --- | --- |
| $pHI_0$ | Magnitude of the prior that the actions of others are generally motivated by harmful intent (HI) toward the self, $p(HI)^{t=0}$. Increasing this parameter increases the belief that a partner is motivated by harmful intent before any actions are observed. |
| $pSI_0$ | Magnitude of the prior that the actions of others are generally motivated by self-interest (SI) irrespective of the self, $p(SI)^{t=0}$. Increasing this parameter increases the belief that a partner is motivated by self-interest before any actions are observed. |
| $uPri$ | Uncertainty over priors. Increasing this parameter broadens the prior distribution of both $p(HI)^{t=0}$ and $p(SI)^{t=0}$. |
| Prior | $p(HI)^{t=0} \sim Bin(HI; pHI_0, uPri, NB)$ $p(SI)^{t=0} \sim Bin(SI; pSI_0, uPri, NB)$ $p(HI, SI)^{t=0} = p(HI)^{t=0} p(SI)^{t=0}$ $NB = 9$ |
| $w_0$ | Intercept of the likelihood matrix, $\pi_{gen}$, that calibrates the magnitude of attributional change when a fair or unfair action is made by a partner. |
| $w_{HI}$ | Impact on beliefs that an outcome (r) is motivated by harmful intent. Increasing this parameter leads to greater influence of a partner's behaviour on attributions of harmful intent (belief flexibility). |
| $w_{SI}$ | Impact on beliefs that an outcome (r) is motivated by self-interest. Increasing this parameter leads to greater influence of a partner's behaviour on attributions of self-interest (belief flexibility). |
| Likelihood | $\pi_{gen}(r = 0; HI, SI) = \sigma(w_0 + [w_{HI} * HI - \delta] + [w_{SI} * SI - \delta])$ $\pi_{gen}(r = 0.5; HI, SI) = 1 - \pi_{gen}(r = 0; HI, SI)$ $\delta = \frac{NB + 1}{2}$ $\sigma(x) = \frac{1}{1 + e^{-x}}$ |
| Update | $p(\widehat{HI}, \widehat{SI})^t = \frac{\pi_{gen}(r; HI, SI) p(HI, SI)^{t-1}}{\sum_{HI', SI'} \pi_{gen}(r; HI', SI') p(HI', SI')^{t-1}}$ |
| $u\pi$ | The consistency with which partners were believed to act in accordance with their character. Higher values reduce consistency, causing a partner's behaviour to have less impact on beliefs. |
| Consistency rule | $p(HI, SI)^t \propto p(\widehat{HI}, \widehat{SI})^t \frac{1}{u\pi} + \xi$ $\xi = 0.02/NB^2$ |
| $\eta$ | Controls the mixture of prior and posterior beliefs used as a starting point for each new encounter. Higher values indicate more reliance on information gathered from the last encounter, rather than reverting to prior beliefs. The product from the below equation, $\overline{p(HI, SI)}^{t=C}$ replaces $p(HI, SI)^{t-1}$ when beginning a new encounter. |
| Change point | $\overline{p(HI, SI)}^{t=C} = p(HI, SI)^{t=0} * [1 - \eta] + p(HI, SI)^{t=C} * \eta$ $C = \text{final action of an other in an interaction}$ |

The winning model uses eight parameters that calibrate an agent’s initial and ongoing beliefs about others. It encodes the agent’s prior expectations of harm, pHI_0_, and self-interest, pSI_0_, and the certainty of these expectations, uPri. Three parameters implement the agent’s internal likelihood of a partner acting with self-interest or harm based on their behaviour, influencing belief updates (w_0_, w_HI_, w_SI_). A noise parameter (uπ) indicates the agent’s uncertainty over the representation of their partner. The model also includes a belief persistence parameter, η, for agents to either persist with their most recent beliefs or re-set them to the prior expectations (above) upon encountering new partners, with higher values indicating less resetting. See table 1 for further details.

All computational models were fitted using a Hierarchical Bayesian Inference (HBI) algorithm which allows hierarchical parameter estimation while assuming random effects for group and individual model responsibility [38]. This process is shown to be most robust to outliers versus non-hierarchical inference or standard hierarchical inference with fixed effects, and minimises parameter and model confusion [38]. Parameters were estimated using the HBI in native space drawing from broad priors (*μ*_*m*_=0, *σ*_*m*_ = 6.5; where *m*={*m*_1,_ *m*_2_, *m*_3_}). This process was run independently for each drug condition due to the dependency of observations between conditions (the same participants were in each condition). Parameters were transformed into model-relevant space for analysis. All models and hierarchical fitting was implemented in Matlab (Version R2022B). All other analyses were conducted in R (version 4.2.3; x86_64 build) running on Mac OS (Ventura 13.0). All statistics are reported as: (X, 95%CI: Y, Z), where X is the regression coefficient, and Y and Z are the 95% lower and upper confidence intervals (CI), respectively. All dependent regressors were centred and scaled. To consider the uncertainty of estimates we conducted Bayesian paired sample t-tests to assess individual-level parameter changes. This used JAGS as a backend MCMC sampler [39]; differences in the mean are additionally reported with their corresponding effect sizes (Cohen’s d) and posterior 95%HDI (High Density Interval). The raw output of this is listed in Table S1. Bayesian paired sample t-tests were also used to assess differences between attributional coupling over time. To note, in the original behavioural analysis [35] we excluded one extra participant due to their extreme trait psychometric paranoia score (leaving 27 participants), however trait paranoia was not the subject of this analysis, and hierarchical model fitting constrains group behaviour during parameter estimation. Nevertheless, for transparency, we include analytic estimates with the original 27 individual included for comparison. This did not change conclusions (Table S2).

We also sought to examine model covariance. Exploratory factor analysis used oblique rotation, including all parameter estimates for each individual within placebo and haloperidol conditions. Optimal factors were determined from observation of the scree plot and cross-validated model accuracy (Figure S9). Cross-validation used 10 folds with three repeats within a logistic general linear model. Parameter loadings and individual factor scores >|0.4| were retained for analysis.

## Results

### Behavioural results

Behavioural results were published previously [35]. To summarise, when averaged over all Dictators, haloperidol caused a reduction in harmful intent attributions versus placebo (−0.17, 95%CI: -0.28, -0.05), but L-DOPA did not. Haloperidol also increased self-interest attributions versus placebo (0.16, 95%CI: 0.05, 0.27), but L-DOPA did not. Unfair and partially fair Dictators both elicited higher harmful intent (Partially fair = 0.28, 95%CI: 0.16, 0.40; Unfair = 0.75, 95%CI: 0.63, 0.87) and self-interest attributions (Partially fair = 0.59, 95%CI: 0.63, 0.87; Unfair = 1.16, 95%CI: 1.05, 1.27) versus fair Dictators.

### Model comparison and recovery

Bayesian hierarchical fitting and comparison identified that at the group level (Figure 2A), participants under placebo and haloperidol were best fitted by model 3. This model assumed agents use a single uncertainty over both attributional priors, although used separate likelihood weights to update their beliefs about their partners’ policy. In contrast, participants under L-DOPA were best fit by model 2. This model assumes participants hold individual uncertainties over their prior beliefs, although use the same likelihood weight to update both attributional dimensions. Importantly, model parameters under L-DOPA were not opposing haloperidol changes vs. placebo, supporting behavioural analyses (see Figure S10).

**Figure 2.**
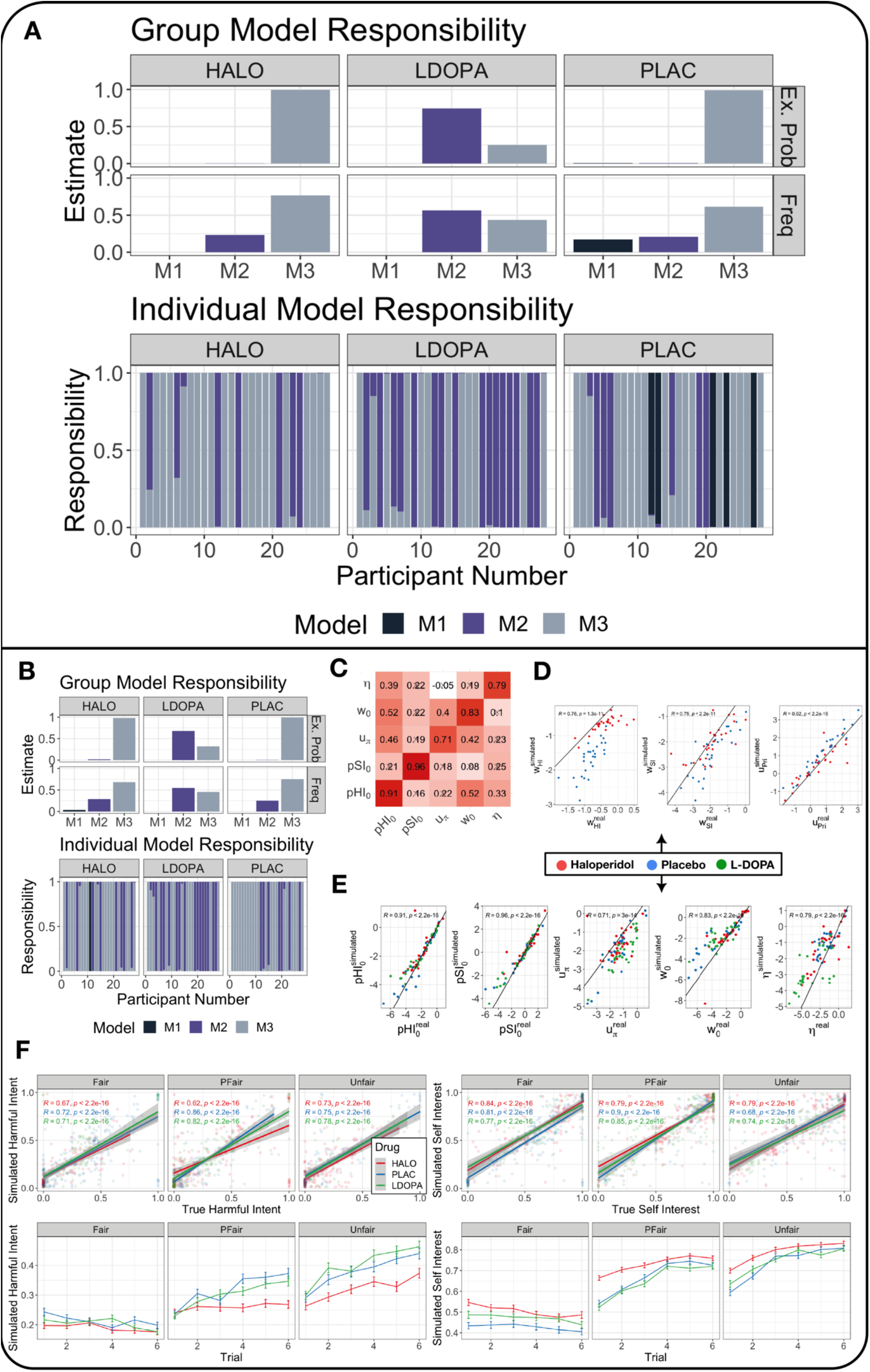
Model comparison, recovery, and generative performance. (A) Model responsibility across all three drug conditions. Greater model responsibility at the group and individual level indicates that a particular formulation was the most likely generative model to explain the data. Ex. Prob = Exceedance probability that a single model best defines group behaviour. Freq = Model frequency that each model is the best fitting model for participants. (B) Model recovery. All recovery analyses used n=28 synthetic participants – one for each real parameter set approximated from the data. The HBI algorithm correctly identified the correct model for most participants with trivial differences between model frequencies. (C) Correlation matrix of common parameters across all drug conditions for simulated (y axis) and real (x axis) data. All correlations were over 0.71 (p values < 0.001). ‘X’ indicates a non-significant association. (D) Individual correlations between common parameters across haloperidol and placebo conditions for simulated (y axis) and real (x axis) data. All correlations were over 0.71 (p values < 0.001). Black lines indicate the linear model of perfect association (*r*=1). (E) Individual correlations between common parameters across all drug conditions for simulated (y axis) and real (x axis) data. Black lines indicate the linear model of perfect association (*r* =1). (F) Top panel: Correlation between simulated and real harmful intent (left) and self-interest (right) attributions across all Dictator policies. Bottom panel: Simulated harmful intent (left) and self-interest (right) attributions for each drug condition and Dictator policy.

For each condition we examined model generative performance and reliability. We extracted parameters for each individual under each condition according to the model that bore most responsibility for their behaviour (Figure 2B). We then simulated data for each participant with their individual-level parameters for each condition and model and re-estimated model comparison, recovered each model, generated attributions for each trial and dictator condition, and fitted regression models for main effects. Bayesian hierarchical fitting and comparison on simulated data demonstrated excellent similarity to group and individual level model responsibility and exceedance probabilities from real data (Figure S1A). Likewise, individual level parameters demonstrated excellent recovery (all Pearson *r* values > 0.71, p values ∼ 0; Figure S1B, C & D). Simulated and real attributions demonstrated excellent recovery across all drug and dictator conditions (all Pearson *r* values > 0.62, p values ∼ 0; Figure S1E). Simulated attributions also recovered the main effects of drug and dictator condition on attributional dynamics: haloperidol demonstrated reductions in harmful intent versus placebo (−0.26, 95%CI: -0.36, -0.16), but L-DOPA did not, and haloperidol increased self-interest attributions versus placebo (0.26, 95%CI: 0.15, 0.37), but L-DOPA did not.

We were most interested in examining the effect of haloperidol versus placebo in order to understand the mechanism behind the observed descriptive behavioural results. As model 3 achieved group-level dominance across both placebo and haloperidol conditions we were able to directly compare all individual-level, winning model parameters between-conditions {*pHI*_0_, *pSI*_0_, *uPri, uπ, η, w*_0_, *w*_H*I*_, *w*_SI_} (Table 1; see below).

### Haloperidol reduces the influence of priors and the precision of harmful intent

We examined the differences between individual level parameters within-subjects for haloperidol versus placebo (Figure 3A; see Figure S4 [Supplementary Materials] for effect sizes). This suggested that haloperidol increased reliance on learning about a partner just encountered, relative to pre-existing prior beliefs about partners in general (*η*; mean diff. = 0.15, 95%HDI: 0.03, 0.26; effect size = 0.66, 95%HDI: 0.22, 1.10). Haloperidol did not influence the consistency with which partners were believed to act in accordance with their character (*uπ*).

**Figure 3.**
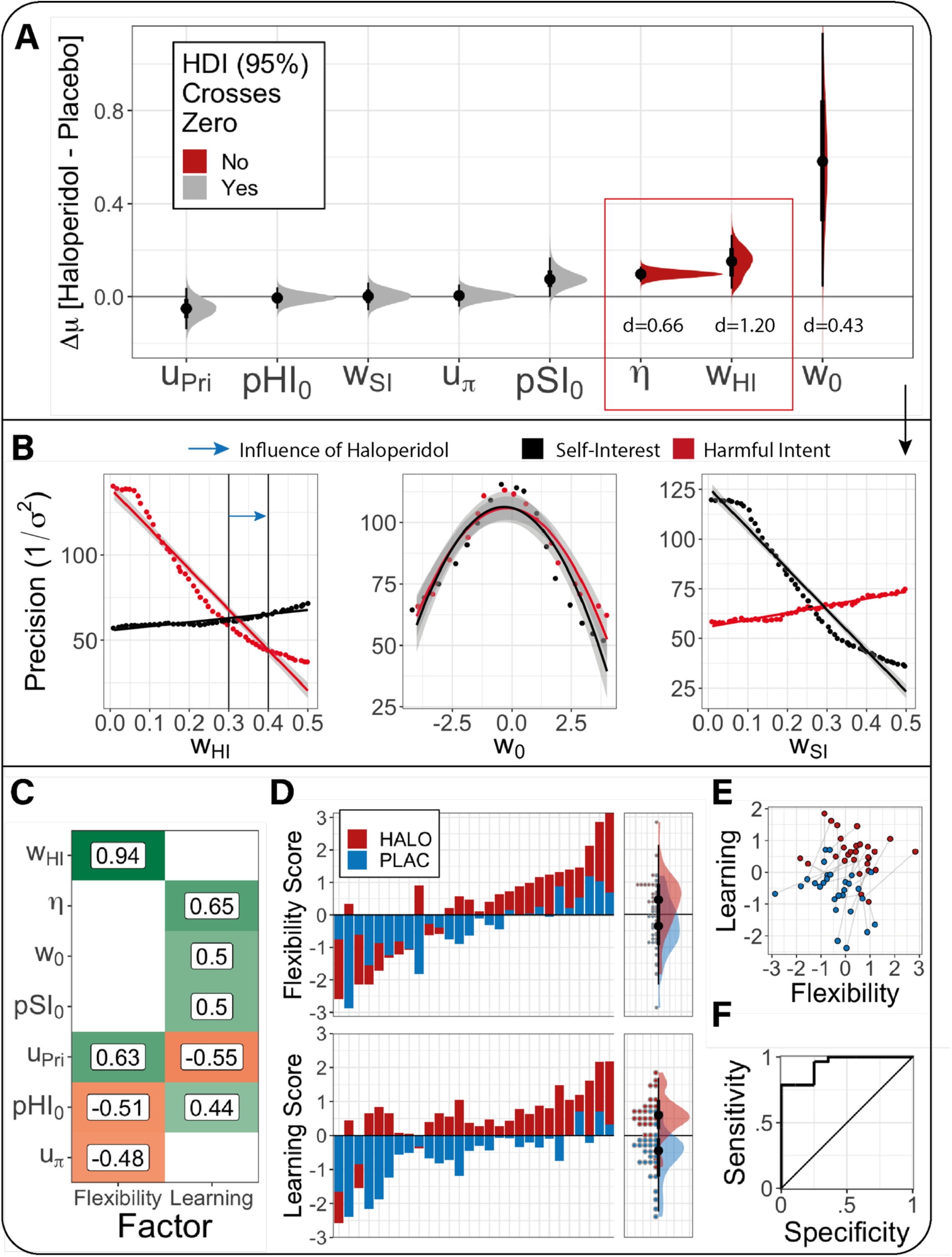
Influence of haloperidol on the winning model. (A) Bayesian t-test results in assessing the difference and uncertainty (distribution of values) of the change in mean (Δ*μ*) in parameter estimates between placebo and haloperidol. Red distributions indicate that the High-Density Interval (HDI) of the mean difference in distributions do not cross 0, suggesting reasonable certainty that the mean difference is not an artefact of statistical noise. ‘d’ values indicate the median effect size (Cohen’s d) for each mean difference (See Figure S4 for distributions). The red box indicates parameters where effect size distributions were most robust, where the 95%HDI and lay outside of the region of probable equivalence with the null hypothesis. (B) Simulations of the marginal effect of likelihood parameters on the precision (1/σ^2^; inverse variance) of harmful intent (red) and self-interest (black) attributions over all trials, controlling for Dictator style. Vertical lines are indicative of the median individual parameter estimates from both haloperidol and placebo groups, with the blue arrow indicating the difference from placebo to haloperidol. For trial-wise and within-Dictator precision changes see Figure S3; to note, simulations are consistent with the notion that *w*_*HI*_ increases flexibility within and between contexts, accentuating smooth learning. To note, there was no significant correlation between *w*_0_, *w*_*SI*_, and *w*_*HI*_ in our parameter estimation from our real data (ps > 0.05; Figure S2) suggesting independent contributions of each to attributional dynamics. (C) Factor loading of each parameter on flexibility (factor 1) and learning (factor 2) dimensions. A loading filter of |0.4| was applied. Both of these factors were able to discriminate most effectively between drug conditions. *w*_*SI*_ is not featured in this plot as it was not meaningfully loaded onto either factor. (D) Factor scores for each individual participant (n=28) for both haloperidol (red) and placebo (blue) conditions ordered from low to high factor loading. The panels on the right of each graph demonstrate the marginal loading across participants. (E) Candyfloss plot of joint factor scores for each individual participant. Grey lines indicate that the same participant was responsible for each connected point under placebo (blue) and haloperidol (red) (F) Receiver Operating Characteristic curve describing the sensitivity and specificity of the combination of flexibility and learning factors on differentiating drug conditions. Area Under the Curve = 0.91. Sensitivity = 0.8. Specificity = 0.78.

Haloperidol increased learning flexibility over harmful intent attributions only. Haloperidol increased the impact of partner behaviour on harmful intent attributions (*w*_*HI*_; mean diff. = 0.10, 95%HDI: 0.06, 0.13; effect size = 1.20, 95%HDI: 0.64, 1.75), but not over self-interest (*w*_*SI*_); a partner’s actions had more impact on a participant’s beliefs about their true motivations of intentional harm. Haloperidol also caused the intercept of the policy matrix to be drawn toward 0, allowing greater updating parity for each unfair or fair partner action (*w*_0_; mean diff. = 0.58, 95%HDI: 0.01, 1.10; effect size = 0.43, 95%HDI: 0.02, 0.82). The *w*_0_ effect size should be treated with caution; the posterior distribution is within the region of practical equivalence (Figure S4).

We sought to further probe the model-based implications of drug differences on attributional flexibility in detail. Simulations on the marginal effect of *w*_*HI*_ on attributional dynamics are suggestive of its role in modulating the precision (1/σ^2^; inverse variance) of attributions over all trials, irrespective of Dictator policy (Figure 3B). To establish this we used a regression model including *w*_*HI*_ as a linear term and *w*_0_ as a quadratic term – this was most parsimonious compared to using *w*_0_ as a linear term (AIC = 568 vs. 1123). There was a main effect of *w*_*HI*_ on the precision of harmful intent attributions (−6.13, 95%CI: -6.28, -5.97; effect size = -0.88, 95%CI: -0.92, -0.85). There was a small effect of *w*_0_ within the same model (−0.06, 95%CI: -0.064, -0.056, effect size = -0.11, 95%CI: -0.14, -0.08). There was a significant but small interaction of *w*_0_ and *w*_*HI*_ on the precision of harmful intent (−0.22, 95%CI: -0.25, -0.20; effect size = -0.05, -0.08, -0.02). Importantly, increased *w*_*HI*_ reduced harmful intent attributions (−0.93, 95%CI: -0.95, -0.92; effect size = -0.13, 95%CI: -0.14, -0.13) through reductions in the precision of harmful intent.

We found evidence that a greater *w*_*HI*_ (cf. effect of haloperidol) may reduce precision most under conditions of ambiguity. Specifically, the precision of harmful intent attributions is lower in partially fair vs fair Dictators (−0.24, -0.33, -0.15; effect size = -0.24, 95%CI: -0.33, -0.15), but unfair vs fair Dictators produced equivalent precision. Dictator policy interacts with *w*_*HI*_: higher *w*_*HI*_ is associated with lower precision under partially fair vs. fair dictators (−0.77, 95%CI: -1.42, -0.42; effect size = -0.11, 95%CI: -0.21, -0.02). Thus, higher *w*_*HI*_ accentuates flexibility within and between partners, but most in ambiguous social contexts where paranoia often flourishes. There was no interaction for unfair dictators vs. fair dictators (Figure S5).

Haloperidol had no net significant influence on *pHI*_0_, *uPri*, or *pSI*_0_ (see Table S1). Individual parameter analysis suggests that haloperidol has a predominant net influence on the flexibility of belief updating about a specific context, here, that of our task. Under the influence of haloperidol, participants’ assumptions about each new encounter are more amenable to change under the influence of recent encounters.

### Alterations to single parameters drive model covariation that differentiates haloperidol from placebo

From our analysis we can conclude that the model is accounting for the true observed data relatively well. Isolated parameter changes between conditions suggest this effect is primarily driven by increases in the impact of partner behaviour on beliefs about harmful intent, *w*_*HI*_, and increased learning from experience, *η*. Considered separately, these key parameters did not fully explain how the model accounted for behaviour changes induced by haloperidol (Figure S4). We therefore sought to identify, through exploratory factor analysis, meaningful patterns over the covariation induced by Haloperidol.

We found that three factors best accounted for the data (Figure S9) with the first demonstrating the greatest eigenvalue (factor 1=2.82; factor 2=1.36; factor 3=1.13). K-fold cross-validation within a logistic model demonstrated that a two-factor solution provided the best median accuracy to discriminate between drug condition (mean accuracy = 0.86) and had the lowest AIC (40.3; see Fig S9). Each factor was able to predict drug condition independently (Factor 1 = 1.52, 95%CI: 0.50, 2.91; Factor 2 = 3.08, 95%CI: 1.72, 5.03), and there was a large effect found between conditions using Bayesian paired t-tests (factor 1: mean diff. = 0.76, 95%HDI = 0.37, 1.17; effect size = 0.94, 95%HDI = 0.35, 1.59; factor 2: mean diff. = 1.34, 95%HDI = 0.87, 1.85; effect size = 1.23, 95%HDI = 0.64, 1.84; Figure 3F).

Factor 1 (Flexibility; Figure 3C) was typified by high values of *w*_*HI*_, and greater consistency between beliefs that a partner’s actions are indicative of their true motivations, *u*_*π*_. Factor 2 (Learning; Figure 3C) comprised high values of *η*, larger intercepts over the policy matrix, *w*_0_, and higher values over priors *pSI*_0_. *pHI*_0_ and *u*_*pri*_ were oppositely loaded onto each factor and would likely nullify each other in cases where participants scored strongly on both (Figure 3E). We note that *pHI*_0_, and *u*_*pri*_ load with slightly more absolute value on the Flexibility factor. For completeness, the third factor was comprised exclusively of *w*_*SI*_ above a cut-off of |0.4| (loading = 0.99), although was not found to be a meaningful factor in differentiating drug scores following cross-validation and logistic model comparison.

### Haloperidol compresses the dimensionality of partner policies

Finally, we explored the impact of haloperidol on attributional coupling: the dependency between intentional attributions over time. This allows analysis into the dependency of different intentional components. To calculate this we estimated Spearman correlations between harmful intent and self-interest for each trial across the sample, controlling for the type of Dictator policy affiliated. This revealed that while harmful intent and self-interest are attributed independently of one another under placebo (mean ρ[sd] = 0.03 [0.07]) replicating prior work [35], under haloperidol they are negatively associated (mean ρ[sd] = -0.22 [0.08]), and this difference is significant (mean diff. = -0.26, 95%CI: -0.32, -0.20; effect size = 2.22, 95%HDI: 1.22, 3.24). This relationship was replicated using simulated model predictions (mean diff. = -0.25, 95%CI: -0.34, -0.17; effect size = -1.53, 95%HDI: -2.28, -0.78); see Figure 4A. There was evidence that the negative association induced under haloperidol decays over time (Pearson *r* = 0.52, p = 0.029). The same is not true under placebo (see Figure 4A). This interaction was not significant (regression coef. = -0.06, 95%CI: -0.12, 0.03).

**Figure 4.**
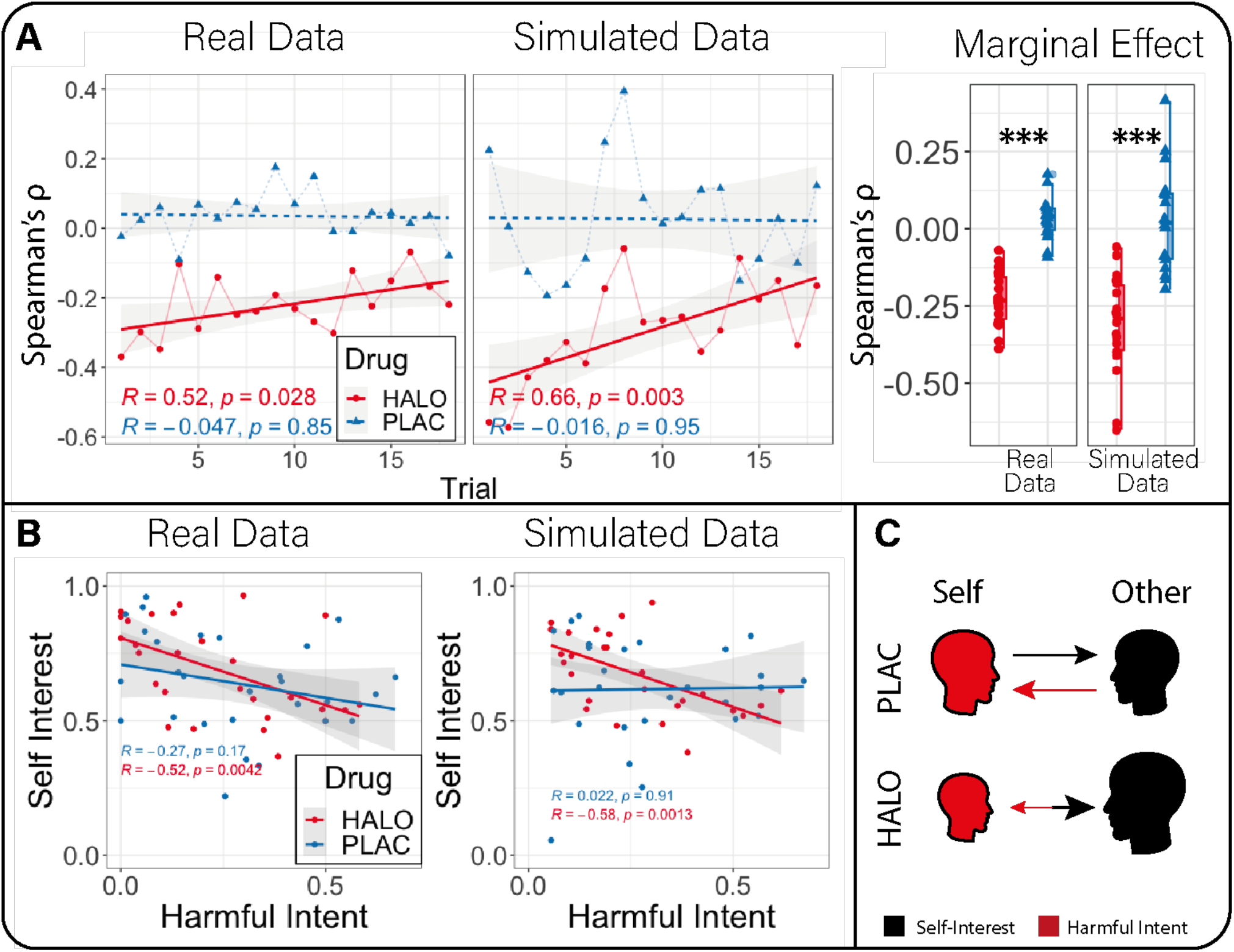
Association of mental state attributions between drug condition. (A) In both real and simulated data, haloperidol (red) versus placebo (blue) induced a trial-wise negative association between harmful intent and self-interest which decayed over time. The right panel shows the marginal effect of trial-wise correlations between conditions. *** = p<0.001. (B) There was a general negative association between harmful intent and self-interest (Pearson correlation) found under haloperidol (red) for average attributions across all 18 trials. This was not true for placebo (blue). (C) Summary of main effects between drug conditions on self and other oriented intentional attributions following social outcomes. Both trial-wise and averaged associative analyses indicate that other-oriented attributions concerning self-interest of others (black) and self-oriented attributions concerning the harmful intent of others (red) are independent under placebo (PLAC) but coupled under haloperidol (HALO). Under haloperidol this coupling is biased toward exaggeration of other-oriented attributions and diminishment of self-oriented attributions.

In sum, haloperidol causes harmful intent and self-interest attributions to become less independent. This means that under haloperidol participants are more likely to believe someone must be more self-interested if they are perceived to be less intentionally harmful.

## Discussion

We sought to identify the computational mechanisms that explain how pharmacological alteration of dopamine function alters attributions of harmful intent, an important feature of paranoia, given our previous findings that haloperidol reduced harmful intent attributions and increased self-interest attributions in healthy participants (see [35] for previously published behavioural analysis). Here, we tested different computational hypotheses to account more mechanistically for these effects. The data were best fit by a model utilising a common uncertainty parameter over priors, but separate likelihood weights for updating attributions. Using this model, we found evidence that haloperidol reduced the precision of harmful intent (but not self-interest) attributions allowing more belief flexibility between partners. Haloperidol also increased the impact of learning from each encounter; participants relied less on their prior beliefs about the population as a whole. These individual parameter effects were embedded within covariational model alterations that together accounted for attributional change under haloperidol. These changes also caused self-interest and harmful-intent attributions to become negatively associated, suggesting a compression of attributions into a single interpersonal dimension under haloperidol. Together our findings indicate haloperidol promotes flexibility regarding attributions of harmful intent to others by reducing the perceived relevance of the actions of others to the self (Figure 5). In clinical environments this may allow space to reframe beliefs.

**Figure 5.**
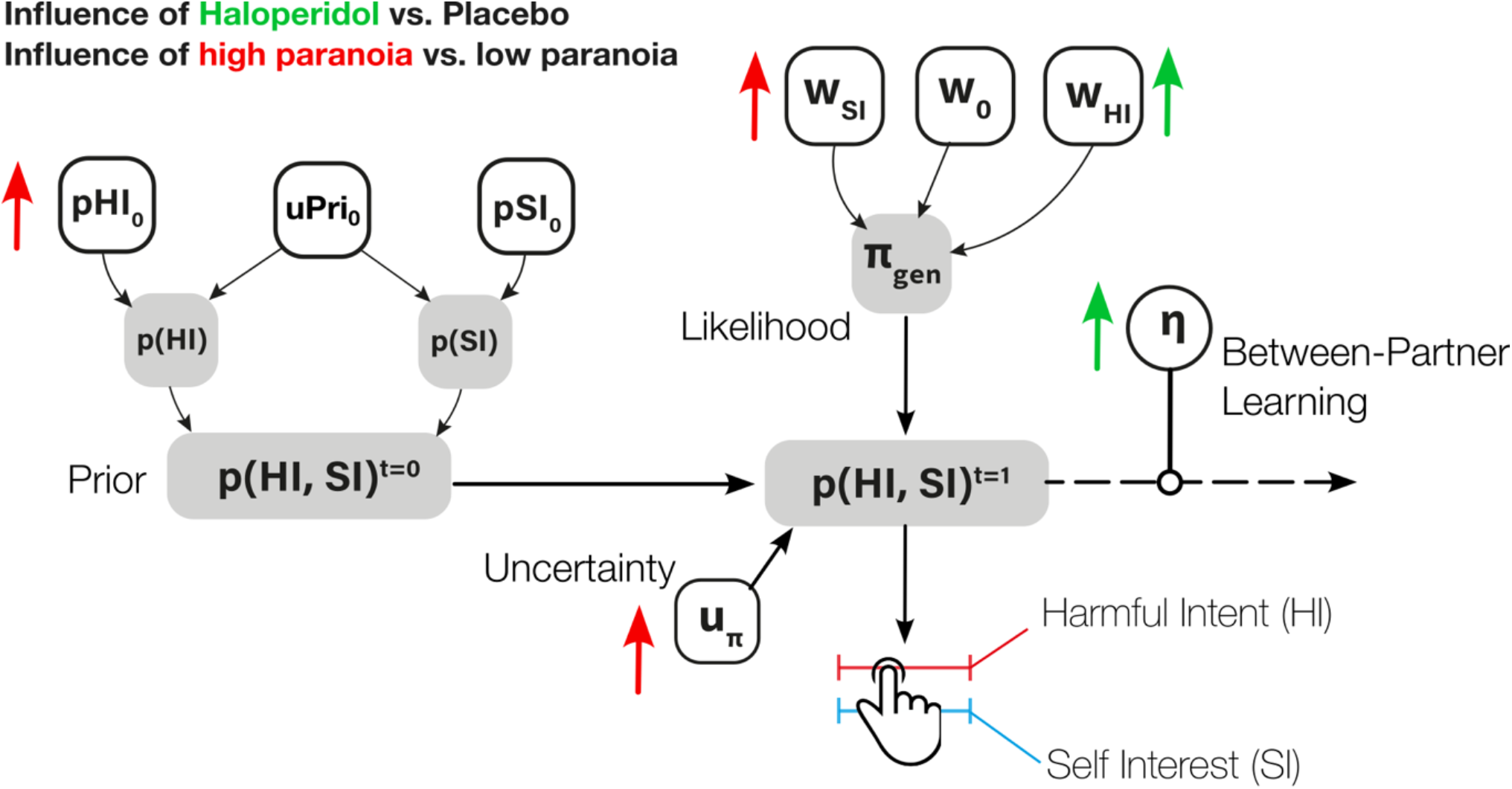
Summary of experimental parameter changes from current and past work. (A) Experimentally observed effects on our model. Overall illustration of the impact of haloperidol on model parameters are illustrated in green. Prior results from the impact of high trait paranoia [30,31] are illustrated in red.

Our findings indicate a reduction in the influence of priors and more flexible beliefs under haloperidol. Previous research links tonic dopamine at D2/D3 receptors to efficient encoding of meaningful stimuli and Bayes optimality [33], cognitive control [40], and sustained attention [41]. Under the model-based, model-free control framework [42], recent work showed D2/D3 antagonism increased model-based control and decision flexibility [21] and increased belief flexibility during a trust game [34]. This may be particularly useful in ‘climbing out’ of paranoia, where one is reluctant to take in positive information about others for fear of ‘false reassurance’. At face value our results conform with previous work: under haloperidol, posteriors are more flexible and less influenced by priors, suggesting more confidence in beliefs about the motivation of partners. However, this general account does not explain why our data show asymmetric decreases in harmful intent and increases in self-interest.

One hypothesis is that haloperidol reduces the perceived self-relevance of outcomes under uncertainty. Social interaction rapidly increases the complexity of possible actions that may be taken. Humans try to reduce this uncertainty by relying on available heuristics, such as using self-preferences as an easily accessible prior belief about others [43-45]. When ambiguity increases, greater uncertainty about others [30,31,19] and environments [20] can increase the perception of social threat. Our analysis suggests that haloperidol may attenuate the relationship between uncertainty and attributions of harmful intent by reducing the perceived self-relevance of others’ actions; attributions of harmful intent, by definition, are inferences about the relevance of threat to the self from another. Given the role of the striatum and medial prefrontal cortex in regulating threat evaluation under stress [46], this reduction in self-relevance may also interact with common neural implementations of self-other modelling [47]; haloperidol may modulate the degree to which information is modelled as self- or other-relevant. The degree to which D2/D3 dopamine receptor function is specific to harmful intent or *all* attributions that are relevant to the self (e.g. altruistic intent of another) can be tested by including an extra dimension within our model; there are a number of hypotheses that can be made with such a modification (see Figure S7).

This pattern leads to a further, complementary proposition: haloperidol may reduce self-relevance through reductions in the complexity or depth of recursive mentalising (how a self thinks about another’s model of the self). In general, the ability to recursively mentalise is computationally expensive [48-50]. Humans try to use cheaper strategies when possible. Recursive mentalising is context dependent: in simple, competitive social scenarios humans are more likely to plan ahead more deeply and entertain recursive beliefs about another’s model of the self [51]. Mentalisation gone awry has also been posited as a core driver of relationship difficulties in clinical populations: paranoia in borderline personality disorder and psychosis are explained as hyper-mentalisation – the inference of overly complex mental states based on sparse data [26,27,52,53]. An alteration in mechanisms that support self-relevant mentalising may explain our findings. This notion is consistent with reported amotivation under haloperidol (individuals are less concerned by outcomes), the role of D2/D3 receptors in promoting cognitive control [40,41], and prior work on the causal role of D2/D3 antagonism in trust behaviours [34]; reductions in the immediate value (and therefore relevance) for the self may facilitate longer term reciprocal trust behaviours without any need to engage deliberate reasoning about future outcomes. A core test of the hypothesis that D2/D3 dopamine is crucial for self-relevant, recursive mentalisation is to use models of hierarchical mentalisation in future experiments that allow estimation of recursive depth in joint social contexts.

The data presented here may be relevant beyond psychiatry. In behavioural economics, there have been several studies on the role of dopamine, reward, and decision making in both social and non-social contexts [54]. Increasing dopamine availability has been shown to increase risky non-social decisions when self-gain is at stake [55], suggesting that dopamine may inflate the attributed value of outcomes to the self. Our data imply that this role of dopamine in modulating monetary value to the self may reflect a broader role in representing the self-relevance of stimuli. The direction of this relationship (self-relevance precedes self-value, or vice versa) is a fruitful target for future research. Our data may also be relevant to the role of dopamine in moral behaviour. In one study, boosting D2/D3 dopamine with pramipexole reduced generosity, especially with close others [56]. Our data complements this work, suggesting that D2/D3 dopamine is involved in calibrating the valuation of self-gain in social decision-making.

On a theoretical level, our formal model distinguishes between computational changes that result from prior representational biases (e.g., higher trait paranoia) and acute state changes during social interaction where potential harm from others is a possibility (Figure 5). Previous modelling with the same task [30] or a reversal variant of the task [31] provided evidence that trait paranoia increases the magnitude of priors over harmful intent, the subsequent increase in the belief that the actions of others are not reflective of their true motivations and a reduced willingness to believe that changes to a partner’s behaviour are motivated by changes to their harmful intent. Naturally, this suggested that prior representations bias how social behaviour is interpreted. On the other hand, the present models suggest that haloperidol acts through increased reliance and impact of likelihoods on the formation of beliefs. Creating phenomenologically plausible formal models that are sensitive to different explanations of behavioural data has been a core aspiration of computational psychiatry [13,14]. Models like ours may be useful in distinguishing between longer term development and near-term alterations in learning that may explain paranoia. Model parameters are constant at the timescale of tasks while potentially evolving at the timescale of personal development, illness and recovery, while learning and inference can be dissected in the timescale of task conditions and trials. Much like prior work distinguishing interventions of representational change (psychotherapy) and emotion modulation (antidepressants; [57]) our model may support similar distinctions following intervention. We thus hypothesise that successful therapeutic use of haloperidol in paranoia will be associated with large changes in likelihood parameters described above but may leave intact, at least in the short term, prior beliefs about the harmful intent of others. D2/D3 independent processes may underpin ongoing vulnerability and may require further psychosocial learning. In our case, our task may only pick up long term representational (prior) changes following extended pharmacological therapy, or in combination with psychological therapy.

We note some limitations. First, we did not use a patient population which means the extent to which the findings generalise to a population with persecutory delusions, rather than non-delusional paranoia, remains unclear. Likewise, in this first study we only included males to avoid hormonal heterogeneity, which might affect drug response and indeed the precise expression of dopaminergic mechanisms [58]. However, this important limitation must be addressed in future studies with studies powered to examine the computational structure of antipsychotic medication in people of different hormonal status and gender. Second, we did not include any non-social comparator (e.g. model-based decision making or volatile environments) when assessing the role of haloperidol on cognition. This leaves a divide between how dopamine influences non-social cognition and mental state inferences. Prior work suggests some shared variance between more foundational computations (e.g. decision temperature, belief updating) and paranoia [20,31,33]. Replicating the present work with non-social comparators of our social task, e.g. using a slot machine partner, may help understand the relations between formal theories of general decision making and how this is expressed at a recursive and intentional level in the same individuals. Third, we did not use a design that probes how dopamine may facilitate generalisation of social knowledge outside of our game theory task. Prior work has demonstrated that representations about learned partners can pass on from one context to another [48]; once a representation is learned using computationally intensive resources, a cheaper, heuristic model can be used. This relates to the question of whether an associative model of updating may be more efficient once a policy is known, and given our findings, whether haloperidol causes a faster transition. Finally, despite the difference in model responsibility, we did not find any influence of L-DOPA on behaviour. This may be due to an insufficient dose or translation of L-DOPA leading to an increase in dopamine release, or the unspecific postsynaptic binding that may result from any successfully increased dopamine release as a consequence of L-DOPA.

## Acknowledgements

The authors would like to thank Uri Hertz who allowed us to use his graphical avatar illustrations for the Sharing Game.

