## Supplementary Materials for "D2/D3 dopamine supports the precision of mental state inferences and self-relevance of joint social outcomes"

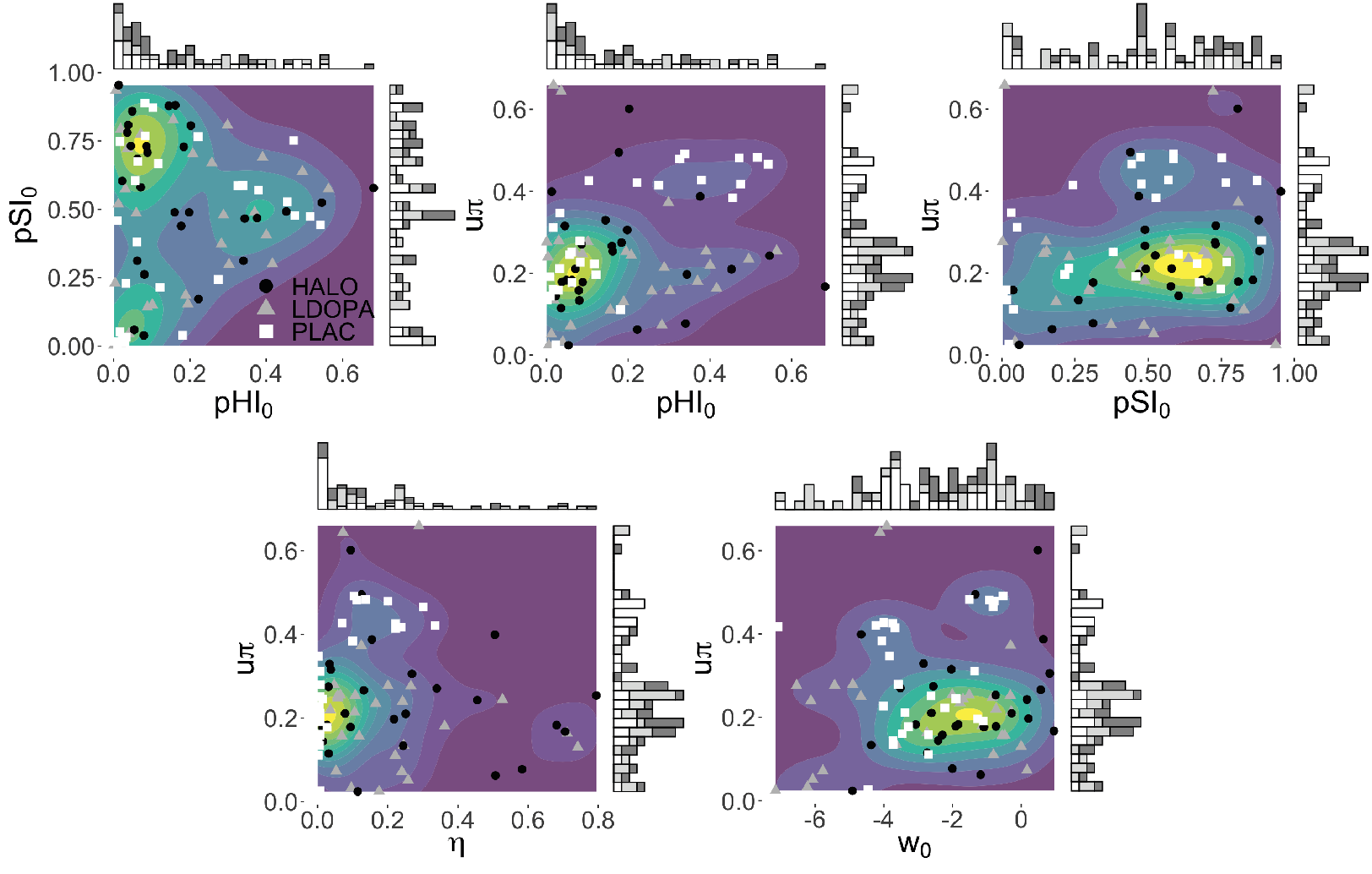
**Figure S1. Common parameter marginal and joint distributions**.

**Figure S2. Common parameter correlations.** Correlations are reflective of the same distributions within figure 2. All correlations coefficients are ranked spearman rho.

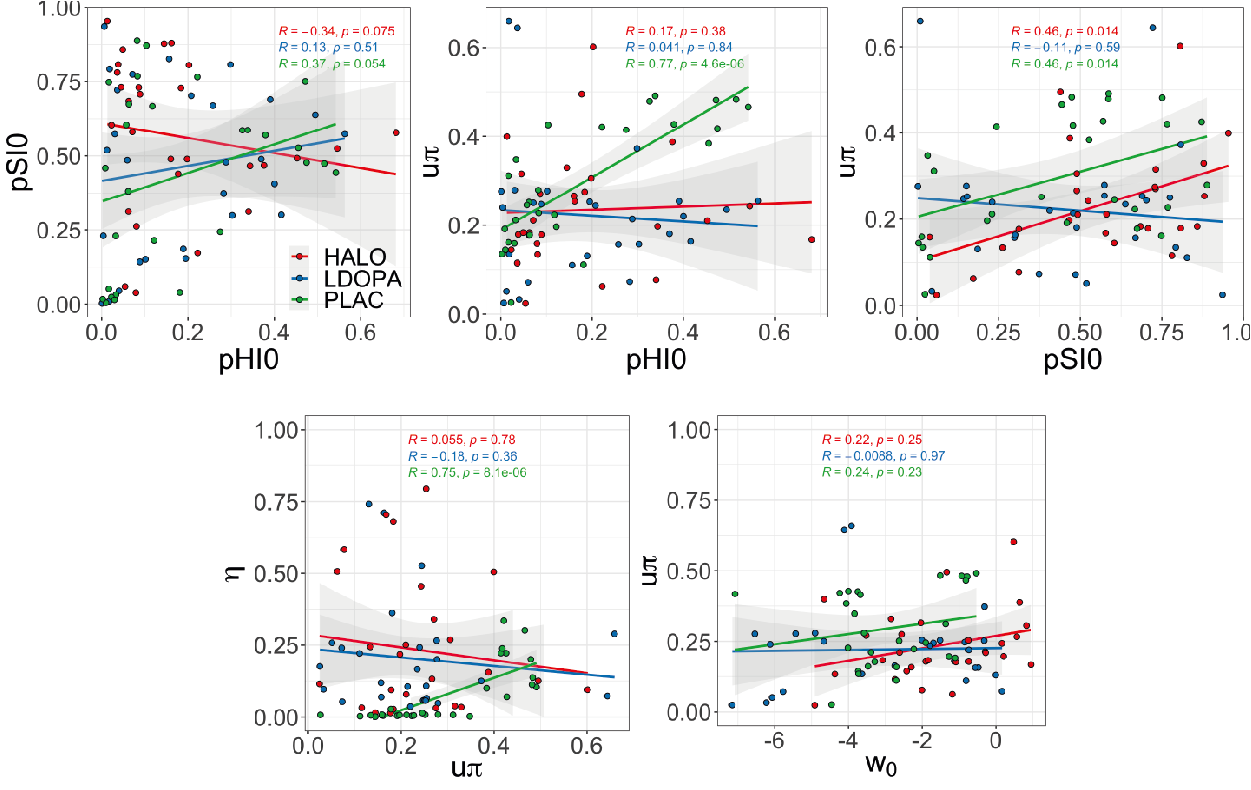

**Figure S3. Parameter correlation matrix (spearman rho) of model 3 parameters under placebo (top) and haloperidol (bottom).** X = not significant.

**Figure S4. Within trial and within-Dictator simulations.** (A) Trial wise marginal effects of each likelihood parameter on harmful intent (red) and self-interest (black) intentional attributions. Simulations control for Dictator policy. (B) Within-Dictator changes in precision as a function of varying $w_{HI}$ values. Marginal influence of Dictator on the precision of harmful intent attributions, irrespective of $w_{HI}$, are 74.6 (fair), 62.8 (partially fair), and 74.1 (unfair). (C) Trial-wise effects of $w_{SI}$ (top) and $w_{HI}$ (bottom) on the magnitude of self-interest and harmful intent attributions, respectively. Increases in each parameter leads to increases in the impact of partner behaviour on attributions.

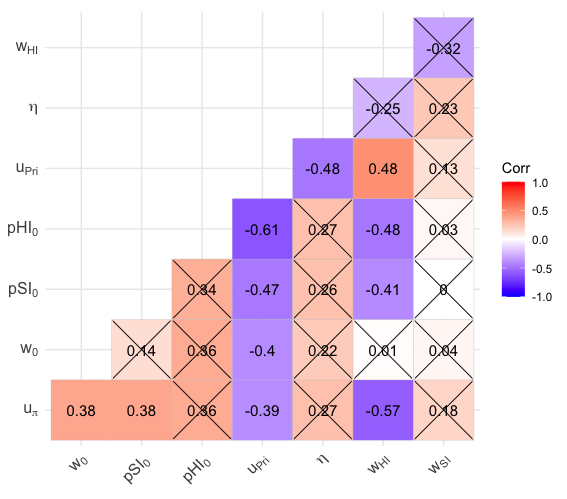

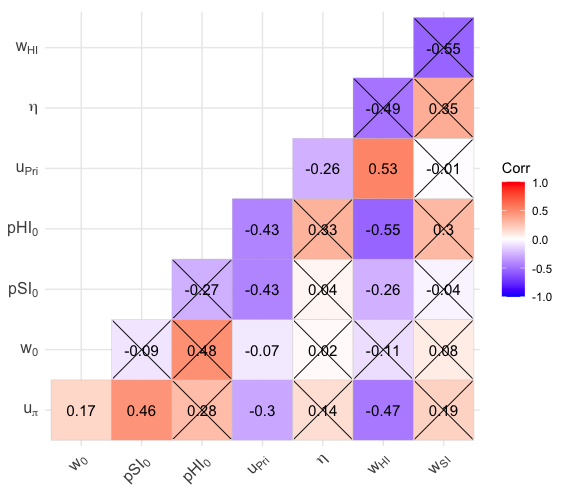

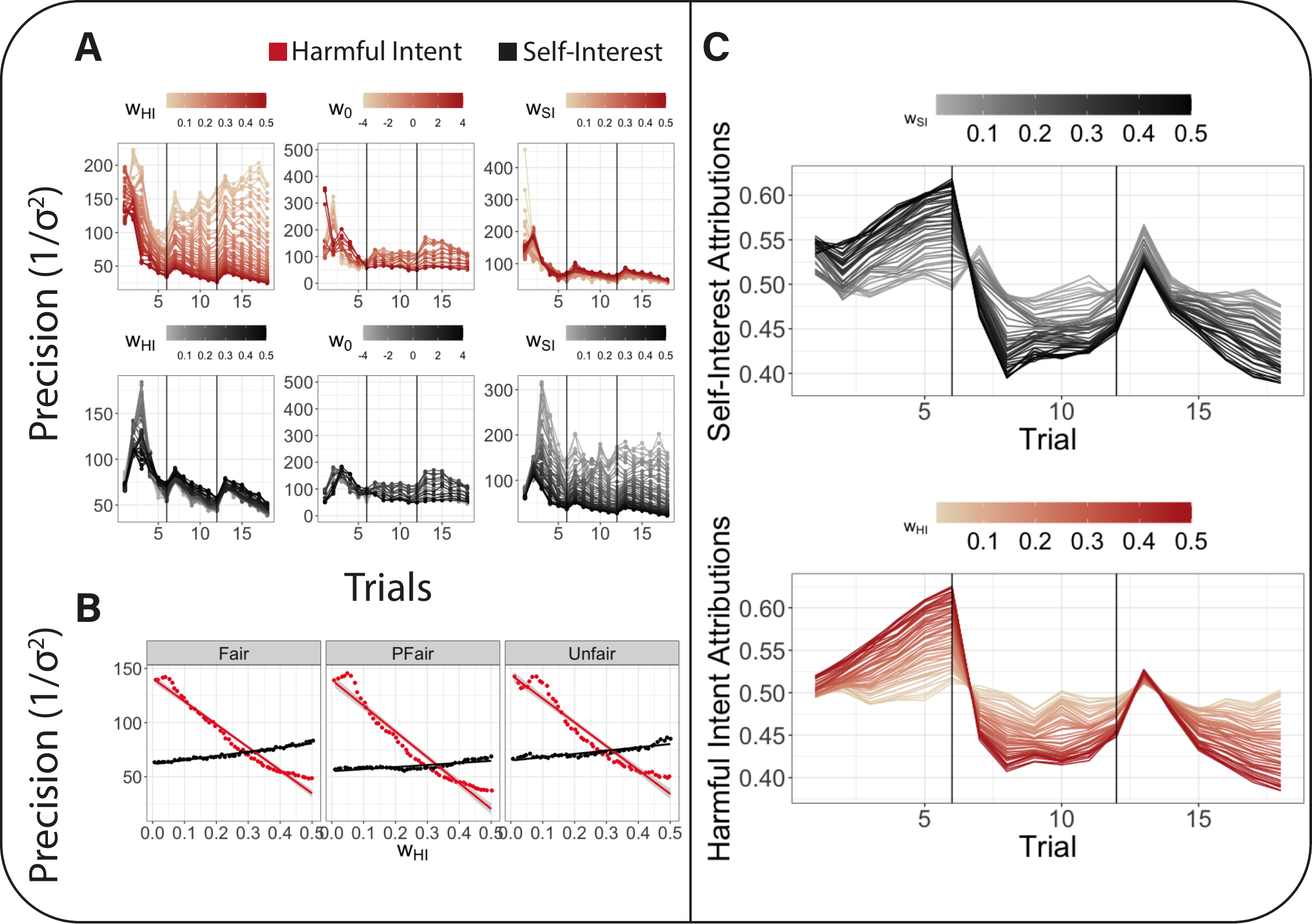

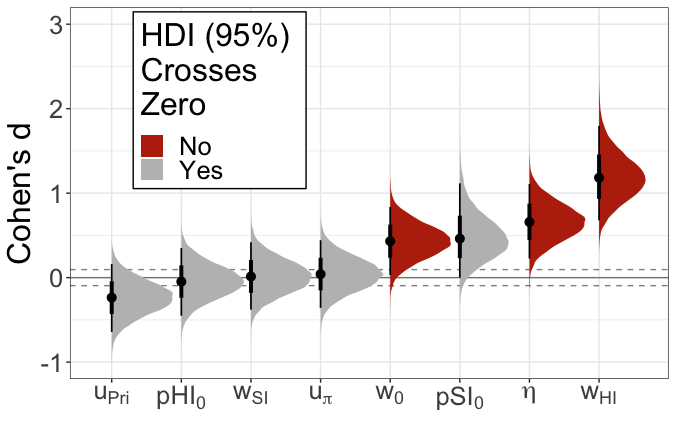
**Figure S5. Effect size posterior distributions from Bayesian paired sample t-tests.** Strongest evidence favoured an influence of haloperidol vs placebo on $w_{HI}$ and $\eta$, with weaker but meaningful effects for$w_{0}$. Red distributions signify that the posterior estimate and 95%HDI does not cross 0. Dotted lines represent -0.095 and 0.095 effect sizes, considered within the region of probable equivalence (ROPE) with the null hypothesis.

**Figure S6. Interaction of** $\boldsymbol{w}_{\boldsymbol{HI}}$ **and** $\boldsymbol{w}_{\boldsymbol{0}}$**, and** $\boldsymbol{w}_{\boldsymbol{SI}}$ **and** $\boldsymbol{w}_{\boldsymbol{0}}$ **on the precision of harmful intent (top) and self-interest (bottom) attributions across contexts.**

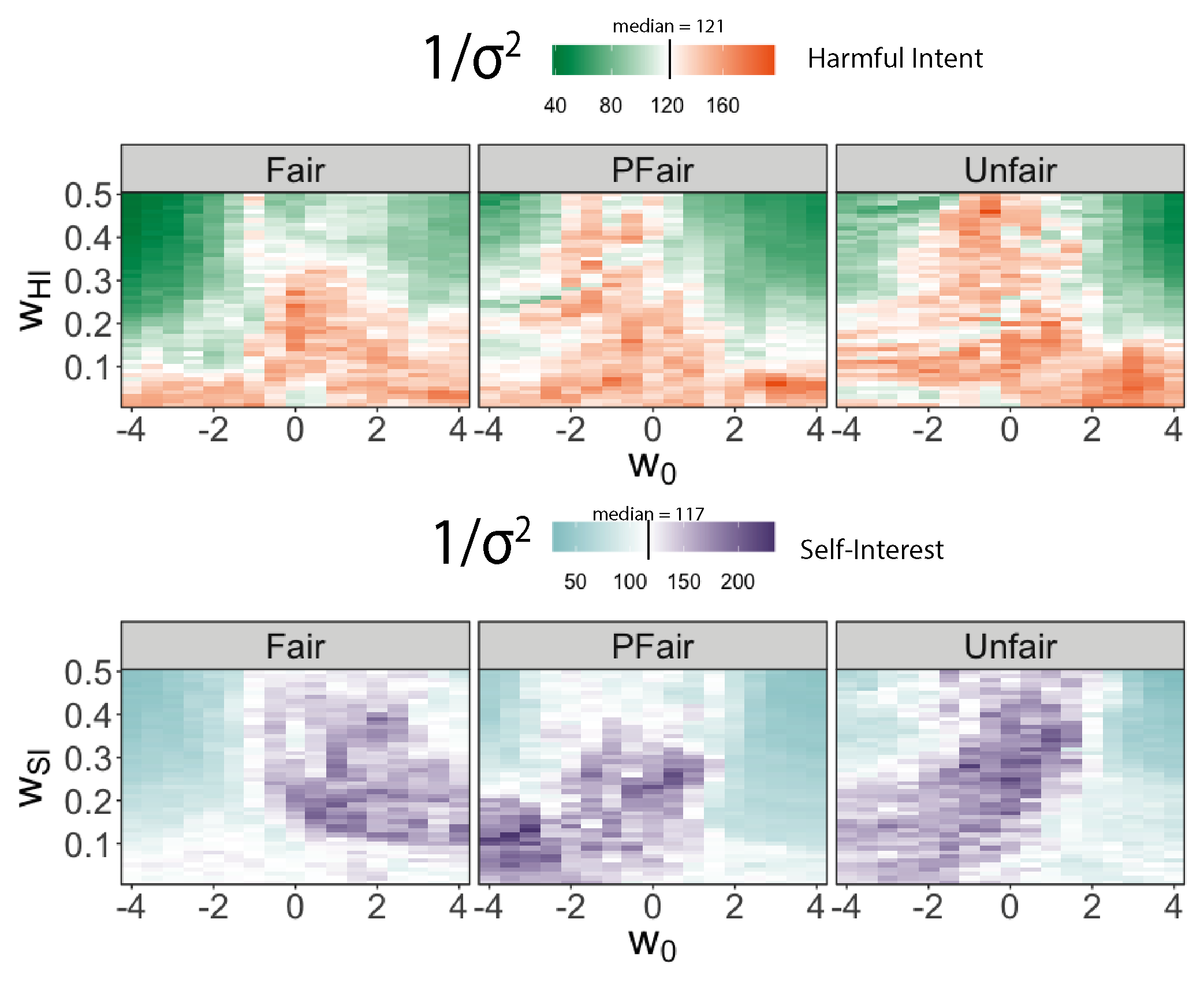

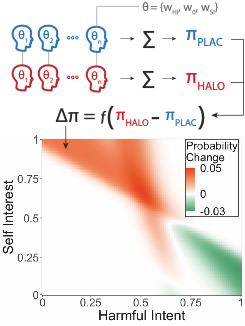
**Figure S7. Full policy matrix difference between PLAC and HALO conditions.** Simulation of the group-level change in unfair action policies. This matrix governs the probability of a partner’s harmful intent (x axis) and self-interest (y axis) given an unfair action. Participant’s individual policy parameters ($w_{0}, w_{HI}, w_{SI})$ for model 3 were extracted and policy maps were simulated, weighted by sample size, and summed for placebo and haloperidol conditions. The summed policy map for placebo was subtracted from the haloperidol policy map to create a difference matrix and was scaled for interpretability. The plot shows the probability of a reduction (green) and increase (orange) in joint attributional probability of a partners intentions given their action under haloperidol.

**Figure S8. Generative predictions about the effects of haloperidol on a model that includes multiple self-relevant beliefs about the motivations of an other**. Hypothesis 1 (H1) states that haloperidol reduces both self-relevant attributions of intent, including altruistic attributions (AI) and harmful intent attributions (HI) versus placebo. Hypothesis 2 (H2) states that haloperidol only reduces self-relevant HI versus placebo. Hypothesis 3 (H3) states that there is an interaction between attributions, such that haloperidol reduces both self-relevant attributions versus placebo, but harmful intent attributions are reduced to a greater degree. In all cases, in line with our data, haloperidol increases self-interest attributions (SI) versus placebo. Such a modification may also be tied with pre-trial predictions of partner behaviour (whether they will be fair or unfair) to further establish a relationship between prediction, outcome, and attribution. HALO = haloperidol; PLAC = placebo.

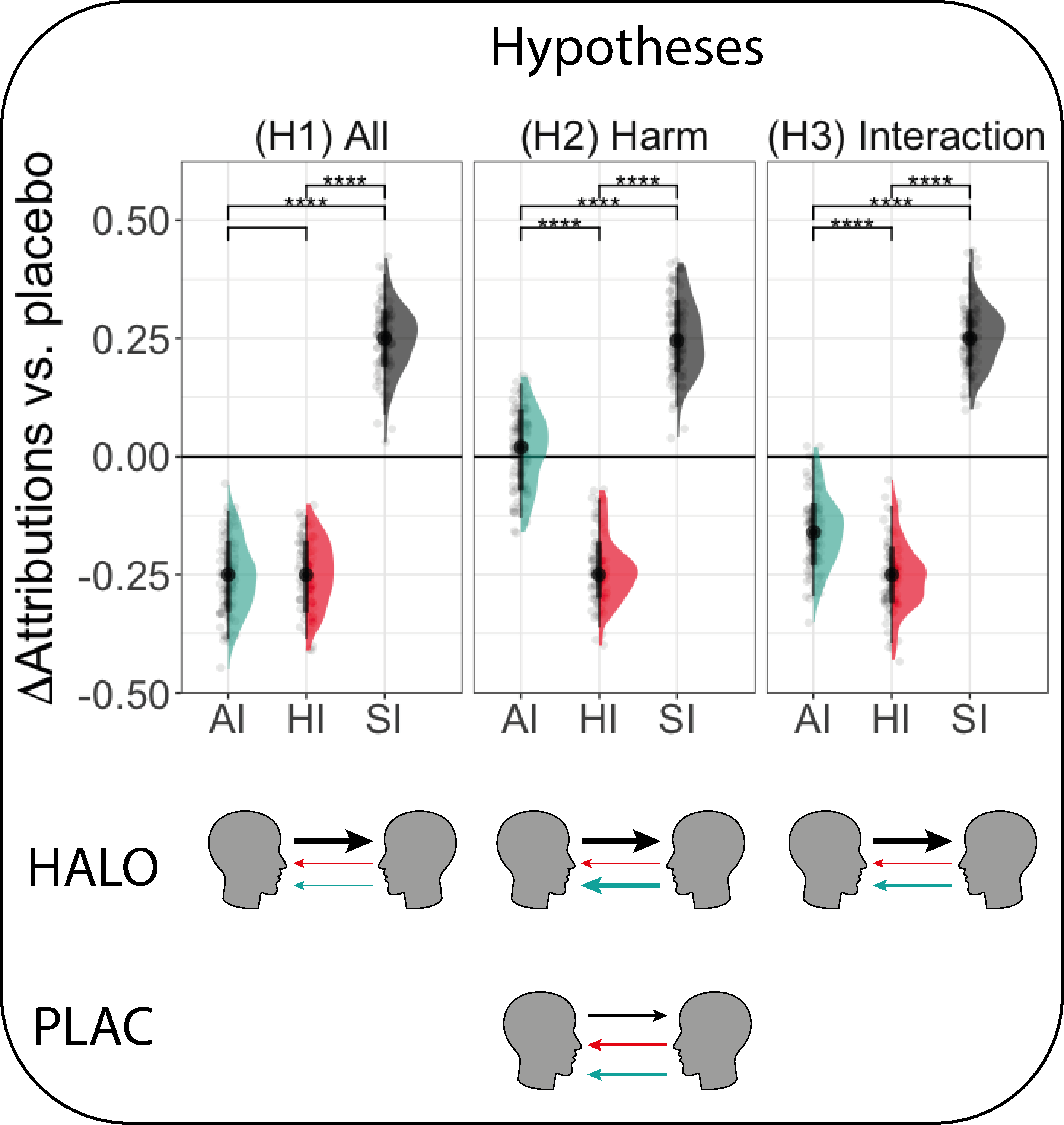

**Figure S9. Scree plot and cross-validated model comparison.** Top panel: scree plot following extraction of eigenvalues of each potential factor loading that may explain the data. Bottom panel: model accuracy for a logistic glm that included only factor 1 (model1), factor 1 and factor 2 (model 2), or all three factors (model 3).

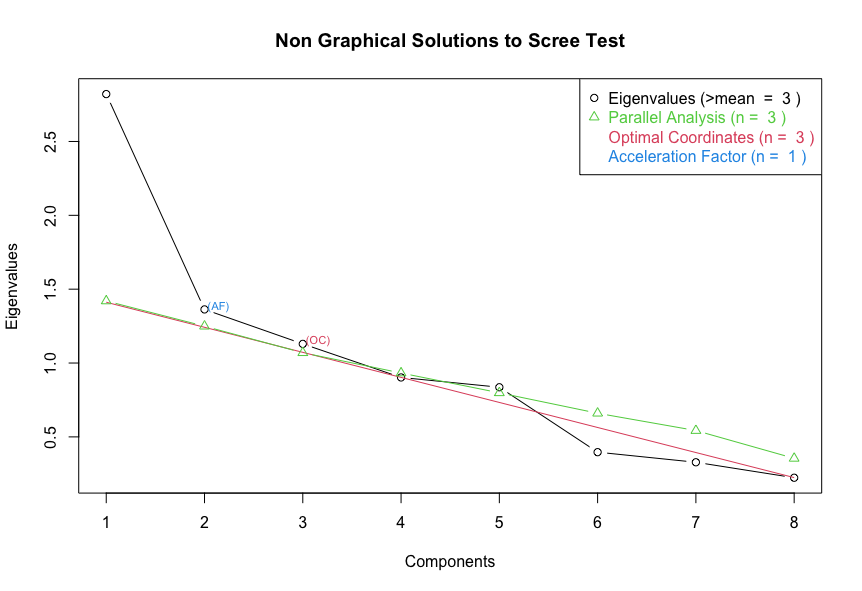

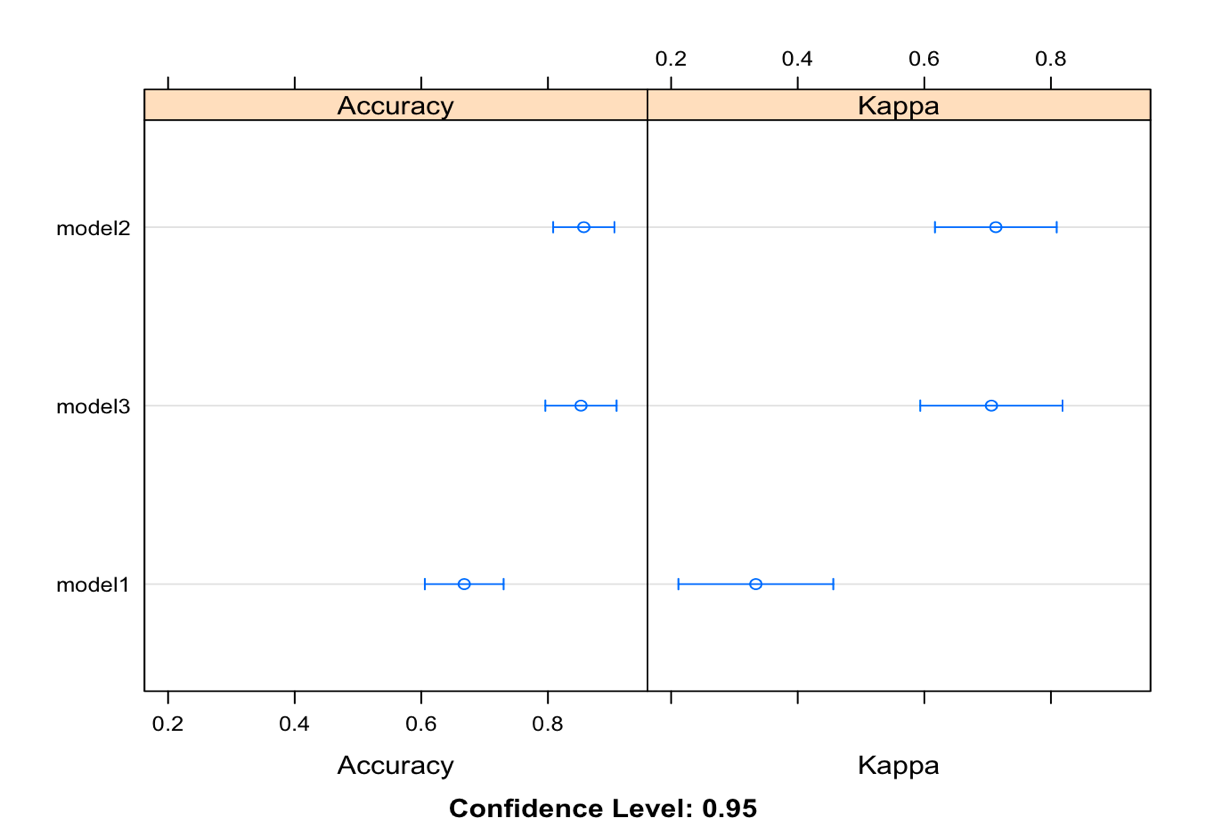

**Figure S10. Relationship between core model parameters for LDOPA and haloperidol.** Values are in native space and taken from the winning model for each condition (Model 2 for LDOPA and Model 3 for haloperidol). Given that LDOPA showed no descriptive overall effect on attributional outcomes, if LDOPA were showing negative associations with haloperidol it would greatly weaken the conclusion that haloperidol was changing model parameters in a meaningful way.

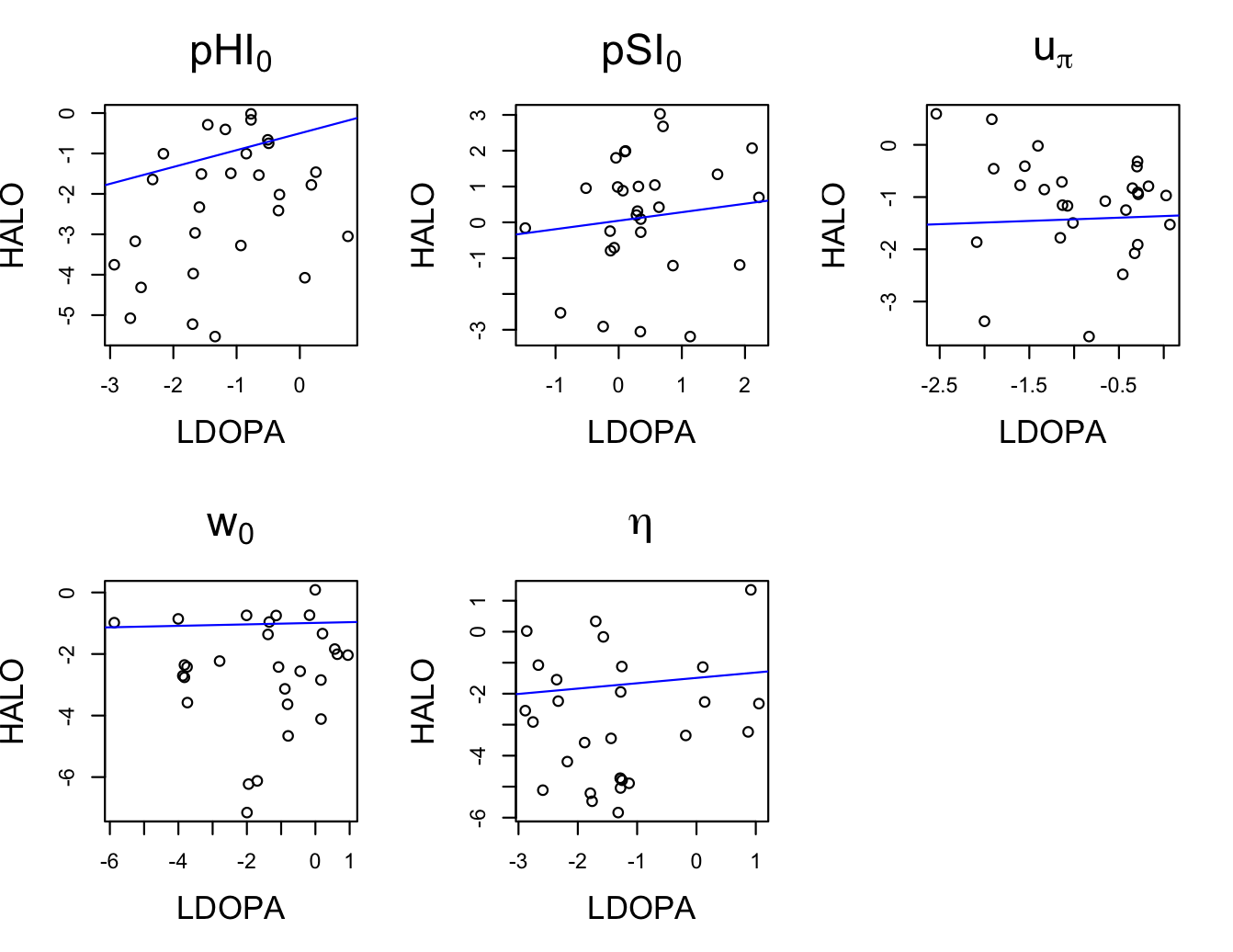

**Table S1. Raw output from Bayesian paired t-tests (n=28).** SD = Standard Deviation, SE = Standard Error. Grey shaded columns = effect size 95%HDI do not cross 0. Effect sizes are in bold. $w_{0}$ posterior effect distributions were within the region of practical equivalence (-0.095, 0.095) and should be treated with caution.

|  | $u_{\pi}$ | η | $p{SI}_{0}$ | $p{HI}_{0}$ | $w_{0}$ | $w_{HI}$ | $w_{SI}$ | $u_{Pri}$ |
| --- | --- | --- | --- | --- | --- | --- | --- | --- |
| Mean Difference | 0.00 | 0.15 | 0.08 | -0.01 | 0.58 | 0.10 | 0.00 | -0.05 |
| SD | 0.02 | 0.06 | 0.04 | 0.02 | 0.28 | 0.02 | 0.03 | 0.04 |
| 95%HDI (lower) | -0.04 | 0.03 | -0.01 | -0.05 | 0.02 | 0.06 | -0.06 | -0.14 |
| 95%HDI (upper) | 0.05 | 0.26 | 0.16 | 0.04 | 1.11 | 0.13 | 0.06 | 0.04 |
| 2.5% | -0.04 | 0.03 | 0.00 | -0.05 | 0.04 | 0.06 | -0.06 | -0.14 |
| 25% | -0.01 | 0.10 | 0.05 | -0.02 | 0.40 | 0.09 | -0.02 | -0.08 |
| Median | 0.00 | 0.15 | 0.07 | -0.01 | 0.58 | 0.10 | 0.00 | -0.05 |
| 75% | 0.02 | 0.19 | 0.10 | 0.01 | 0.76 | 0.11 | 0.02 | -0.02 |
| 97.5% | 0.05 | 0.27 | 0.17 | 0.04 | 1.13 | 0.13 | 0.06 | 0.04 |
| SE | 0.00 | 0.00 | 0.00 | 0.00 | 0.00 | 0.00 | 0.00 | 0.00 |
| Effect Size | 0.04 | **0.66** | 0.48 | -0.05 | **0.43** | **1.20** | 0.02 | -0.24 |
| 95%HDI (lower) | -0.35 | **0.22** | -0.05 | -0.45 | **0.02** | **0.64** | -0.39 | -0.64 |
| 95%HDI (upper) | 0.45 | **1.10** | 1.06 | 0.35 | **0.84** | **1.75** | 0.42 | 0.16 |
| $\hat{R}$ | 1.0008 | 1.0001 | 1.0004 | 1.0001 | 1.0004 | 1.0006 | 1.0004 | 1.0000 |
| n_eff_ | 18767 | 2593 | 9995 | 18133 | 17341 | 19648 | 17702 | 18206 |

**Table S2. Raw output from Bayesian paired t-tests (n=27).** Analysis excludes the participant listed in the original behavioural analysis (Barnby et al., 2020a). SD = Standard Deviation, SE = Standard Error. Grey shaded columns = effect size 95%HDI do not cross 0. Effect sizes are in bold.

|  | $u_{\pi}$ | η | $p{SI}_{0}$ | $p{HI}_{0}$ | $w_{0}$ | $w_{HI}$ | $w_{SI}$ | $u_{Pri}$ |
| --- | --- | --- | --- | --- | --- | --- | --- | --- |
| Mean Difference | 0.00 | 0.16 | 0.08 | -0.01 | 0.55 | 0.10 | 0.00 | -0.05 |
| SD | 0.02 | 0.06 | 0.05 | 0.02 | 0.29 | 0.02 | 0.03 | 0.05 |
| 95%HDI (lower) | -0.05 | 0.03 | -0.01 | -0.06 | -0.03 | 0.06 | -0.07 | -0.14 |
| 95%HDI (upper) | 0.04 | 0.27 | 0.17 | 0.03 | 1.10 | 0.13 | 0.06 | 0.04 |
| 2.5% | -0.05 | 0.04 | 0.00 | -0.06 | -0.01 | 0.07 | -0.07 | -0.14 |
| 25% | -0.02 | 0.11 | 0.05 | -0.03 | 0.36 | 0.09 | -0.02 | -0.08 |
| Median | 0.00 | 0.16 | 0.08 | -0.01 | 0.55 | 0.10 | 0.00 | -0.05 |
| 75% | 0.01 | 0.20 | 0.11 | 0.01 | 0.74 | 0.11 | 0.02 | -0.02 |
| 97.5% | 0.04 | 0.28 | 0.18 | 0.04 | 1.13 | 0.13 | 0.06 | 0.04 |
| SE | 0.00 | 0.00 | 0.00 | 0.00 | 0.00 | 0.00 | 0.00 | 0.00 |
| Effect Size | -0.03 | **0.66** | 0.45 | -0.09 | **0.40** | **1.21** | -0.01 | -0.22 |
| 95%HDI (lower) | -0.42 | **0.21** | -0.06 | -0.49 | **0.00** | **0.65** | -0.40 | -0.63 |
| 95%HDI (upper) | 0.39 | **1.10** | 0.99 | 0.33 | **0.82** | **1.81** | 0.41 | 0.19 |
| $\hat{R}$ | 1.0001 | 1.0017 | 1.0005 | 1.0002 | 1.0003 | 1.0007 | 1.0004 | 1.0006 |
| n_eff_ | 17944 | 3482 | 11273 | 17463 | 17679 | 17634 | 17264 | 18063 |
